# Retrocopy formation and domestication shape genome evolution in sloths and other xenarthrans

**DOI:** 10.1101/2025.09.25.678567

**Authors:** Marcela Uliano-Silva, Helena Beatriz da Conceição, Rafael L. V. Mercuri, Sylke Winkler, Gabriela D. A. Guardia, Eugene Myers, Shane McCarthy, Alan Tracey, Alexander Suh, Mark Blaxter, Pedro A. F. Galante, Camila J. Mazzoni

## Abstract

Xenarthrans, comprising sloths, anteaters, and armadillos, represent one of the most morphologically and physiologically specialised mammalian clades, yet the genomic basis of their adaptations remains poorly understood. Here, we present chromosome-level genomes for the two-toed sloth (*Choloepus didactylus*) and the southern anteater (*Tamandua tetradactyla*), and investigate how retrotransposon-mediated gene duplications (retrocopies) have shaped xenarthran genome evolution. Comparative analyses revealed that xenarthran genomes harbor the highest number of retrocopies reported among mammals, with lineage-specific insertion dynamics. Anteater and armadillo genomes contain older LINE1 repertoires and species-specific older retrocopy insertions. In contrast, sloths retain both an abundance of young LINE1s and thousands of young retrocopies, alongside a large shared set that originated from an evolutionary burst of retroduplication in the branch leading to their last common ancestor (∼30 Mya). In *C. didactylus,* approximately 50% of retrocopies are expressed, compared with 24% in *Dasypus novemcinctus*. Evolutionary analyses identified 38 retrocopies with strong hallmarks of domestication in *C. didactylus*. Many of these retrocopies derive from parental genes involved in mitochondrial and metabolic processes, suggesting a genomic mechanism underlying the physiological specialisations of sloths. Altogether, our findings identify retrotransposition as a major contributor to the genomic architecture of Xenarthra and highlight retrocopy origination as a mechanism for generating lineage-specific novelty and distinctive biological specialisations.

## Introduction

Sloths, anteaters, and armadillos comprise the superorder Xenarthra, one of the four deep-branching clades of placental mammals and the only one to have originated in South America^1,2^. They arose in the Late Paleocene (approximately 60–70 Mya) and once included hundreds of species, from giant ground sloths to glyptodonts and pampatheres, but following terminal Pleistocene mass extinctions, only 43 species remain^3^, distributed in two orders: Cingulata (armadillos) and Pilosa (anteaters and sloths). Xenarthrans display striking morphological and physiological adaptations. Armadillos are the only extant mammals with articulated osteoderms, forming a protective carapace, and have robust, clawed forelimbs adapted for digging and, in some species, conglobation^4,5^. Pilosa includes anteaters, obligatory myrmecophagous with elongated rostra, vestigial or absent teeth, and long, protrusible tongues for feeding on social insects, and sloths (suborder Folivora). Modern tree sloths (*Choloepus* and *Bradypus)* are the only obligatory suspensory quadrupeds among mammals, with elongated limbs, curved claws, and specialised musculature that enable energy-efficient, inverted arboreal locomotion^6^. They are also exceptional in their deviation from the typical mammalian “rule of seven” cervical vertebrae^7^: *Choloepus* usually have 5–7, while *Bradypus* have 8–10. Sloths exhibit the lowest metabolic rates recorded among mammals, often less than half of what is expected for their body size, and show reduced muscle mass, slow digestion, and heterothermy, with body temperatures fluctuating around 5 °C^8,9^. These metabolic adaptations, coupled with their distinctive forelimb morphology, permit a finely tuned and maximally slothful life.

Available xenarthran genomes have limited assembly contiguity, hampering large-scale comparative studies and leaving much of their nuclear genome evolution unexplored, particularly the repetitive elements that comprise substantial portions of mammalian genomes and can be key drivers of genetic innovations. Within this repeat landscape are found retrocopies, gene duplicates generated by the reinsertion of reverse-transcribed copies of spliced, polyadenylated mRNA into the genome^10^. In mammals, retrocopies arise when machinery derived from Long Interspersed Nuclear Element 1 (LINE-1) copies these mRNAs and integrates them via target-primed reverse transcription (TPRT)^11^. The resulting sequences are intronless, often 5′-truncated, and typically lack parental regulatory elements, characteristics that lead these gene duplicates to be classified as processed pseudogenes by some authors. However, even though some retrocopies are not expressed and become pseudogenes, others are “domesticated”. These often show positive selection^10^, are expressed, and can have novel functions as tissue-specific paralogues^12^ or as regulatory noncoding RNAs^13^. These diverse fates mean that retrocopies are both markers of genome history and potential contributors to lineage-specific biological innovations.

Here we present chromosome-level assemblies for the two-toed sloth (*Choloepus didactylus*) and the southern anteater (*Tamandua tetradactyla*), expanding genomic resources for Xenarthra. Using a standardized annotation pipeline, we reveal that xenarthrans harbor the highest number of retrocopies reported in any mammalian group to date, with dynamics that are lineage-specific and closely tied to their LINE1 evolutionary history. In sloths, in addition to a burst of young LINE1s and retrocopies, we identified retrocopies orthologous only within sloths, among which we identified candidates for domestication. Notably, many of these retrocopies are derived from mitochondrial and other metabolic-associated genes, suggesting that these retrotransposition events may contribute to the extreme metabolic adaptations of sloths. Our findings highlight retrotransposition and retrocopies as central forces shaping xenarthran genome architecture, providing a framework for understanding how their dynamics can facilitate the evolution of extreme phenotypes.

## Results

### Two New High Quality Genomes for Xenarthra

We generated genome assemblies for *Choloepus didactylus* and *Tamandua tetradactyla* using the Vertebrate Genomes Project (VGP) v1.6 Assembly Pipeline (see Methods). Both genomes were sequenced using PacBio CLR long reads, scaffolded with Bionano optical maps, chromatin conformation capture (Hi-C) data, and manually curated to achieve high-quality, chromosome-level assemblies. The final assemblies have scaffold N50s of 146 Mb and 174 Mb, respectively (Table 1). The X and Y sex chromosomes were reconstructed in both assemblies. The large majority of the assembled sequences were assigned to chromosomes in both assemblies (99.94% in *C. didactylus* and 99.51% in *T. tetradactyla*; Supplementary Figure 1A).

**Table 1:**
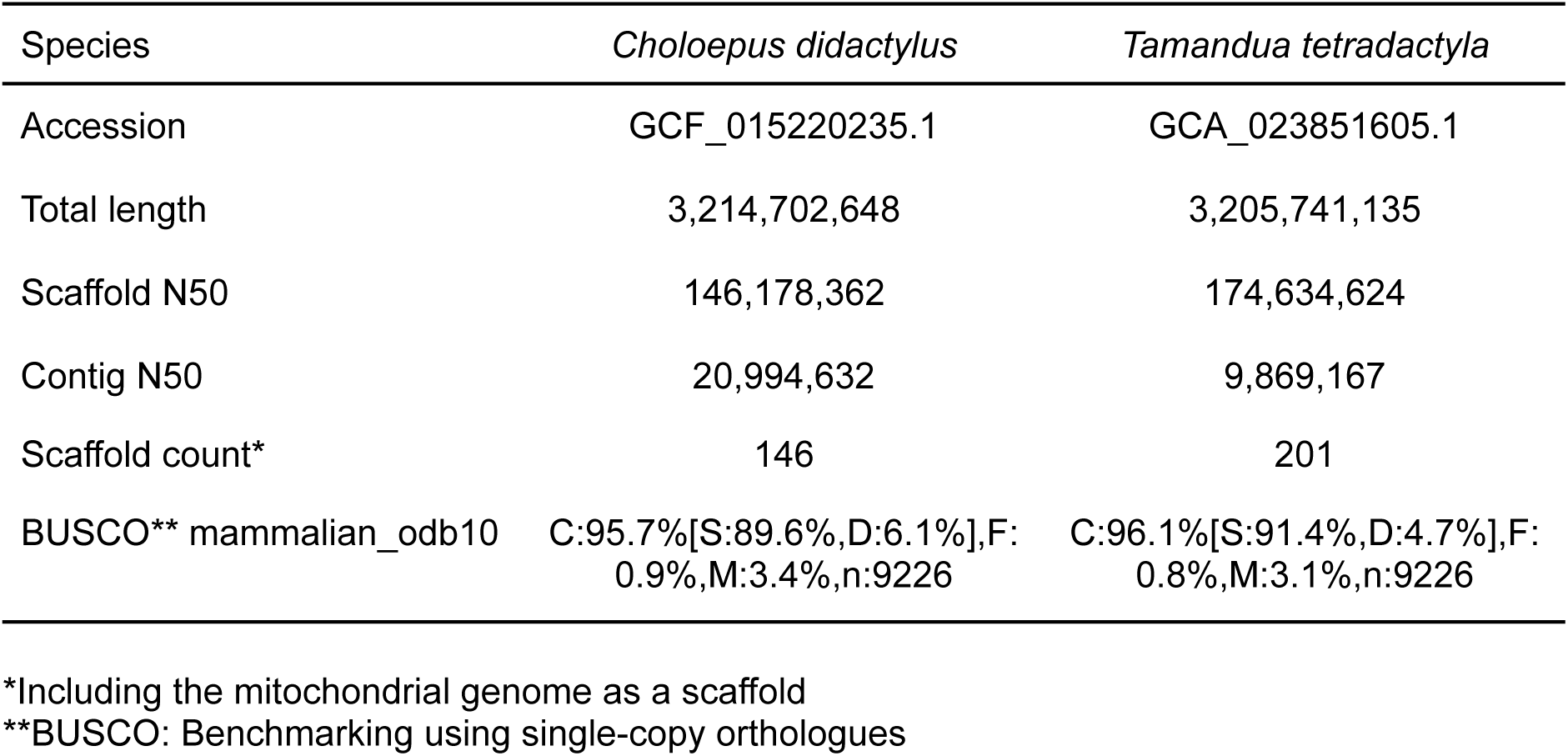
General metrics of *Choloepus didactylus* and *Tamandua tetradactyla* genome assemblies.

To investigate chromosome structure and genome-wide synteny in Xenarthra, we performed synteny analysis using Benchmarking Universal Single-Copy Orthologs (BUSCO^14^) orthologues across five species: the two newly sequenced genomes (*C. didactylus* and *T. tetradactyla*) and three publicly available genomes from the sloths *Bradypus torquatus*^15^ and *Choloepus hoffmanni,* and the armadillo *Dasypus novemcinctus* (Figure 1, Supplementary Table 1). The analysis reveals conserved chromosome architecture in Xenarthra. The karyotype of *T. tetradactyla* appears to be the most divergent among the five species. In contrast, the three sloths and *D. novemcinctus* share more similar karyotypes, suggesting greater chromosomal stability in these lineages (Figure 1).

**Figure 1.**
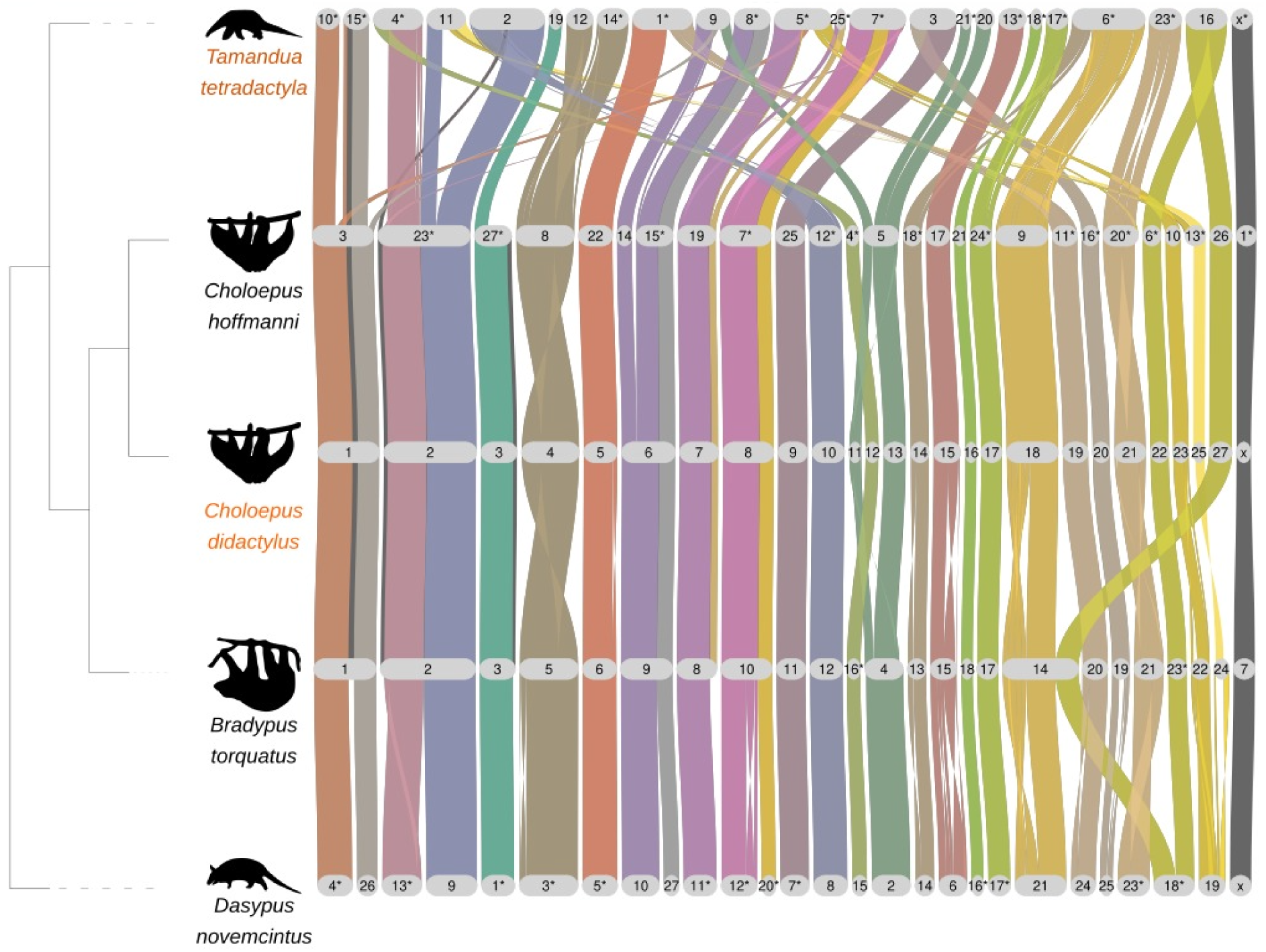
BUSCO-based synteny across xenarthran genomes. BUSCO-predicted genes and conserved syntenic blocks are shown for five Xenarthra genomes, visualised using GENESPACE^16^. The species highlighted in orange (*Choloepus didactylus* and *Tamandua tetradactyla*) are genome assemblies generated in this study. In each species, grey lozenges indicate assembled chromosomal pseudomolecules. Colored ribbons connect orthologous blocks between species, asterisks represent flipped orientation. The cladogram on the left indicates phylogenetic relationships among the species.

### Recent LINE1 activity distinguishes sloth genomes within Xenarthra

To investigate repeat landscape evolution in Xenarthra, we merged species-specific libraries into a common library and used this to annotate all genomes consistently (see Methods). As in most mammals, xenarthran genomes are dominated by LINEs (Figure 2A), with LINE1 being the most abundant family (Supplementary Figure 2). However, in contrast to humans and other xenarthrans, sloth genomes show pronounced peaks of LINEs with low divergence from the inferred consensus, indicating recent bursts of activity (Figure 2A).

**Figure 2.**
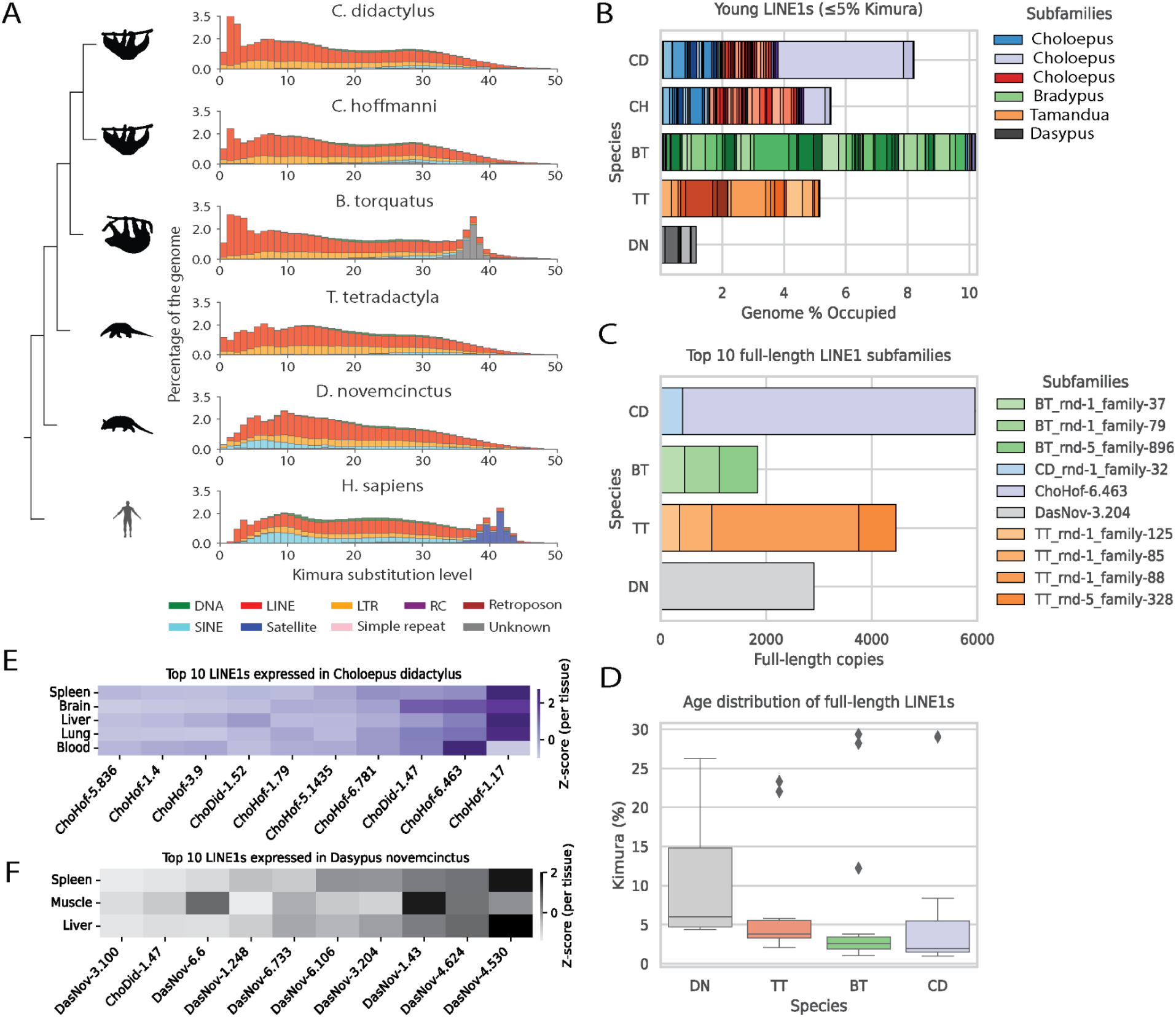
Repeat landscapes in xenarthrans and humans. A) Kimura-corrected repeat landscapes for xenarthran and human genomes aligned to the species cladogram. B) Proportion of the genome occupied by LINE1 subfamilies. C) Total counts of LINE1 full-length copies for the 10 most abundant LINE1 subfamilies. D) Mean Kimura divergence of all LINE1s identified as full-length. E–F) Expression of LINE1 subfamilies in *C. didactylus* (E) and *D. novemcinctus* (F). Species abbreviations: CD, *C. didactylus*; CH, *C. hoffmanni*; BT, *B. torquatus*; TT, *T. tetradactyla*; DN, *D. novemcinctus*.

We examined these young LINE1s in more detail by quantifying the contribution to genome content of subfamilies with a Kimura divergence of ≤5% across species. Sloths have a larger proportion of their genomes occupied by young LINE1s compared with other xenarthrans, with *B. torquatus* showing the highest proportion (Figure 2B). Within sloths, most LINE1 subfamilies are shared between *C. didactylus* and *C. hoffmanni*, consistent with their more recent divergence (∼5 Mya)^17^. In contrast, *B. torquatus* shares only a subset with *Choloepus*, and most of its young LINE1s are species-specific, reflecting its earlier divergence within Folivora (∼30 Mya)^17,18^. Beyond sloths, no young LINE1 subfamilies (≤5% divergence) are shared between all sloths and *T. tetradactyla* or *D. novemcinctus*, both of which harbor mostly species-specific expansions.

We also identified thousands of full-length LINE1s in all xenarthran genomes, except for *C. hoffmanni*, where only nine were recovered (Figure 2C). The highest counts were found in *C. didactylus* (6,700 full-length copies), followed by *T. tetradactyla* (5,159). In most species, expansions were driven by a single dominant, species-specific LINE1 subfamily (Figure 2C). The subfamilies forming full-length LINE1s tend to be younger in sloths than in the anteater and armadillo (Figure 2D). For instance, *C. didactylus* is dominated by 5,531 full-length copies of subfamily ChoHof-6.463 (Kimura ∼2%), while the most abundant family in *T. tetradactyla* is 2,785 copies of TT_rnd-1_family-88 (Kimura ∼2.4%), and in *D. novemcinctus* is 2,910 copies of DasNov-3.204 (Kimura ∼5.2%).

To assess possible LINE1 transcriptional activity, we analyzed RNA-seq data (see Methods) from *C. didactylus* and *D. novemcinctus* (Figures 2E and F). Several LINE1 subfamilies showed detectable expression in both species. In *C. didactylus*, the young ChoHof-6.463 subfamily (Kimura ∼2%, 5,531 full-length copies) had higher expression than most other LINE1s, with particularly elevated levels in blood tissue. In contrast, *D. novemcinctus* expressed DasNov-3.204 (Kimura ∼5.2%, 2,910 full-length copies), but this was not among the most highly expressed LINE1s. Instead, its top-expressed elements (DasNov-4.530, DasNov-4.624, DasNov-1.43) all lack full-length representatives and have mean Kimura divergences >7%, suggesting pervasive or readthrough transcription of fragments rather than autonomous LINE-1 activity in *D. novemcinctus*.

These results indicate that each xenarthran species exhibits a distinct LINE1 dynamic, reflecting their evolutionary divergence. In sloths, however, several lines of evidence point to more recent and potentially ongoing LINE1 activity. These include the high genomic proportion of young LINE1s (Figure 2A, B), the abundance of full-length elements and their younger age distribution (Figure 2C, D), and the elevated expression of a recently expanded subfamily in *C. didactylus* (Figure 2E). This contrasts with the older LINE1 expansions observed in *T. tetradactyla* and *D. novemcinctus*, where ancient insertions predominate and younger elements show small or non-detectable expression.

### Retrocopies are abundant in Xenarthra genomes

Because LINE1s provide the enzymatic machinery that drives retrocopy formation from protein-coding genes^11^, we next investigated the abundance and evolutionary dynamics of retrocopies across Xenarthra. Using our standardized annotation pipeline^19^, we catalogued retrocopies in all available xenarthran genomes, including our new *C. didactylus* and *T. tetradactyla* assemblies, and compared them to patterns across 41 mammalian species in our database (Figure 3; https://www.rcpediadb.org/). We broadly define retrocopies as intronless, mRNA-derived inserts, regardless of sequence decay, and reserve the term “putatively domesticated” for the subset that, upon downstream analyses, exhibits hallmarks consistent with neofunctionalisation.

**Figure 3:**
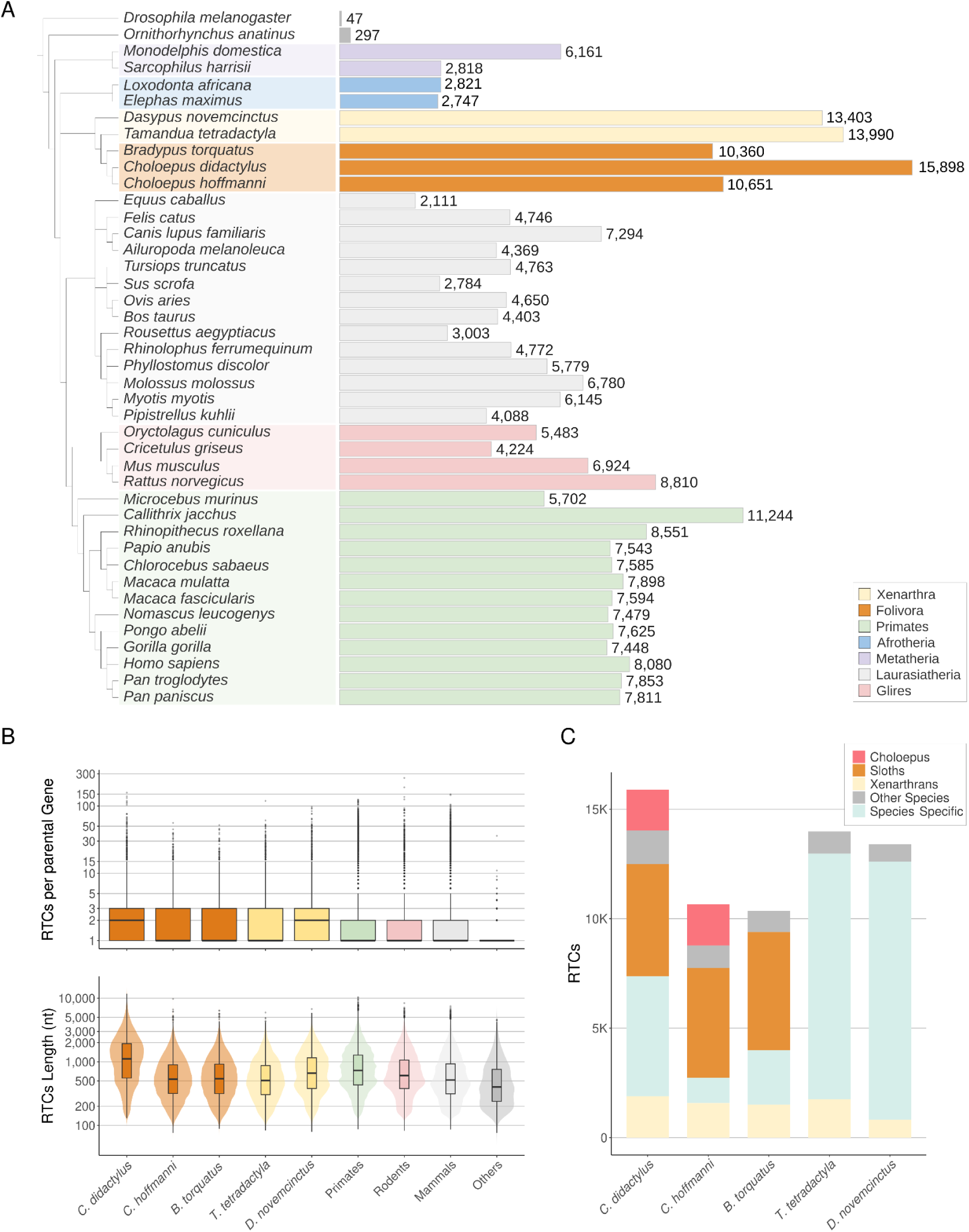
Xenarthra have the highest number of retrocopies among mammals. A) Total retrocopy (RTCs) counts across mammals and *Drosophila melanogaster*. Horizontal bars represent the number of retrocopies identified per species. The phylogenetic tree highlights Xenarthra in yellow and the suborder Folivora (sloths) in orange. B) Retrocopy profiles. Upper panel: box plots showing the number of retrocopies per protein-coding parental gene. Lower panel: violin plots and boxplots showing retrocopy length distributions (in nucleotides). C: Orthology relationships of retrocopies across Xenarthra. Bar heights represent total retrocopy counts per species, with colors indicating orthology groups. ‘Other species’ represent retrocopies with orthology to other mammals.

Xenarthran genomes harbor the highest retrocopy numbers reported in mammals to date (Figure 3A). We identified 13,403 retrocopies in *D. novemcinctus*, 13,990 in *T. tetradactyla*, 10,360 in *B. torquatus*, 10,651 in *C. hoffmanni*, and 15,898 in *C. didactylus*—the latter representing the largest retrocopy count reported for any mammal, or animal, so far. These counts far exceed those in other mammals, which typically range between 2,000 and 8,000 retrocopies per genome^19^, revealing a clear phylogenetic signature of retrocopy expansion in Xenarthra (Figure 3A). The distribution of retrocopies per parental gene follows a consistent pattern across species (Figure 3B, upper panel). In most mammals, parental genes typically generate a single retrocopy, with only a minority undergoing extensive retroduplication^19^. In *C. didactylus* and *D. novemcinctus*, however, the median is two retrocopies per parental gene, and the number of highly duplicated genes is particularly pronounced in *C. didactylus*, with some parental genes giving rise to more than 100 retrocopies. *C. didactylus* also exhibits a distinctive retrocopy length profile (Figure 3B, lower panel). While most species show median retrocopy lengths of 500–750 nucleotides (nt), *C. didactylus* retrocopies are significantly longer, with a median of 1,100 nt (Wilcoxon test, p < 0.0005).

Considering the exacerbated number of retrocopies in Xenarthra, we next explored their evolutionary origins by extracting retrocopy sequences with flanking regions and aligning them across multiple mammalian genomes to assign orthology (see Methods). Only a small fraction (<15% across all species) represents orthologous retrocopies shared among all xenarthrans (Figure 3C). Most retrocopies in *T. tetradactyla* and *D. novemcinctus* are shown as species-specific, and may reflect independent insertion events at either species or clade levels (suborder Vermilingua and order Cingulata, respectively). With three representative species from both genera, we show that sloths retain a large shared subset: over half of retrocopies in *B. torquatus* (52%), and ∼47% and 32% in *C. hoffmanni* and *C. didactylus*, respectively, are exclusive to sloths, consistent with origin in the branch leading to their last common ancestor ∼30 Mya^17,18^, after their divergence from other xenarthrans. Additional retrocopies are shared only between *C. didactylus* and *C. hoffmanni*, consistent with retrotransposition activity in their more recent common ancestor (∼5 Mya). Among sloths, *C. didactylus* shows the largest fraction of species-specific retrocopies (34%), followed by *B. torquatus* (23%) and *C. hoffmanni* (10%).

In addition, to investigate retrocopy age and evolutionary trajectories, we predicted open reading frames (ORFs) and estimated synonymous substitution rates (dS) from codon-aware alignments of retrocopy–parental gene pairs. We then compared dS distributions for species-specific retrocopies and sloth-only orthologs (Figure 4A). As expected, sloth-only retrocopies show no peak near dS = 0, consistent with insertion events predating the divergence of sloths (∼30 Mya) and subsequent sequence decay. By contrast, species-specific retrocopies reveal distinct lineage-specific signatures: in sloths, a substantial fraction of retrocopies fall at dS ≈ 0 (17.6% in *C. hoffmanni*, 15.8% in *C. didactylus*, and 4.6% in *B. torquatus*), indicating recent or ongoing retrotransposition. In *T. tetradactyla* and *D. novemcinctus*, no equivalent peak is observed (Figure 4A), suggesting that their bursts of retrocopy formation occurred earlier and that subsequent divergence has eroded retrocopy sequences.

**Figure 4.**
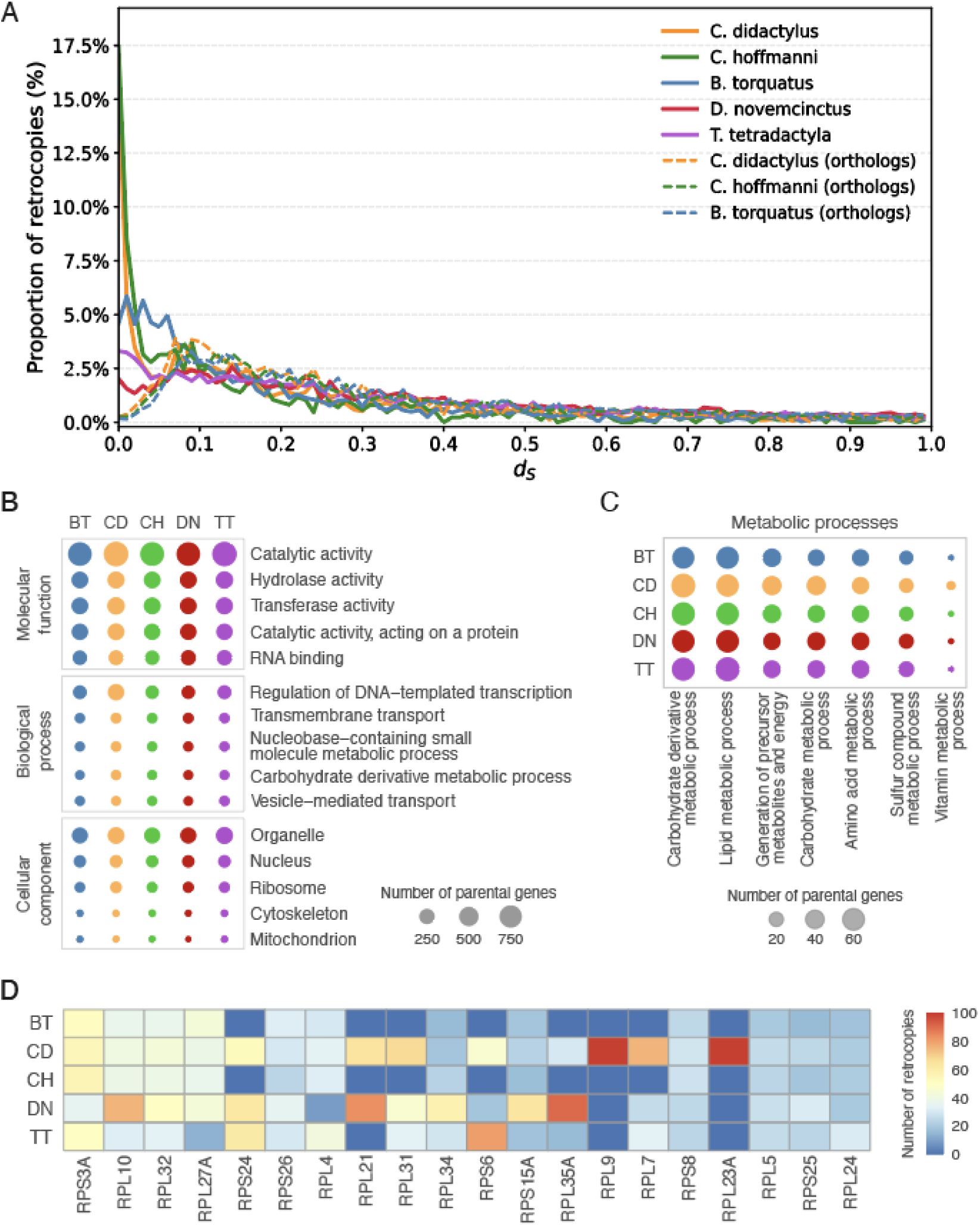
Evolution and function of retrocopies in sloths. **A**: Distribution of synonymous substitution rates (dS) between retrocopies and their parental genes in *B. torquatus*, *C. hoffmanni*, *C. didactylus T. tetradactyla* and *D. novemcinctus*. Solid lines represent all species-specific retrocopies; dashed lines indicate only those found to be orthologous across the three sloth species. Peaks at low dS values, particularly in *C. didactylus, C. hoffmanni* suggest a larger proportion of very recent or preserved retrocopies. **B** and **C:** Most enriched GO Slim terms among parental genes of retrocopies, shared across xenarthran species. Bubble size represents the number of retrocopied parental genes annotated with each term. **D**: counts of the most retrocopied parental genes. Species abbreviations: CD, *C. didactylus*; CH, *C. hoffmanni*; BT, *B. torquatus*; TT, *T. tetradactyla*; DN, *D. novemcinctus*.

Additional analyses of retrocopy-predicted ORFs further highlight these differences, revealing a greater proportion of young and potentially coding retrocopies in *C. didactylus*. This species exhibits the highest fraction of retrocopies retaining ORFs (75.6%), compared with *B. torquatus* (63.2%), *C. hoffmanni* (62.3%), and less than 60% in *T. tetradactyla* and *D. novemcinctus* (Supplementary Figure 3A). *C. didactylus* also harbors the largest absolute number of ORFs (Supplementary Figure 3B). While ORF length distributions are broadly similar across species (medians around 98 amino acids), *C. didactylus* shows a marked skew toward longer ORFs (Supplementary Figure 3C), consistent with more recent insertions and/or improved preservation of coding potential.

Taking these observations together, sloths stand out within Xenarthra for harboring both an enrichment of full-length, low-divergence LINE1 families and peaks of young retrocopies (dS ≈ 0). In contrast, *D. novemcinctus* and *T. tetradactyla* show marks of older LINE1 repertoires and species-specific retrocopies that do not show comparable young dS peaks. These contrasting patterns suggest that retrocopy dynamics in Xenarthra have been strongly shaped by lineage-specific differences in LINE1 activity.

### Functional annotation of parental genes yielding retrocopies

We functionally annotated the parental genes that give rise to retrocopies using Gene Ontology (GO) Slim and found that approximately half of them carry GO annotations in each species. Across Xenarthra, we identified up to 116 distinct GO slim terms, of which 100 were shared among all species, underscoring the strong convergence in the functional landscape of retrocopy-generating genes. The most represented categories included catalytic, transferase, and hydrolase activities (reflecting core metabolic processes) together with more general roles, such as retrocopies of parental from ribosomal proteins, nucleic acid binding, and transcriptional regulation genes (Figure 4B). These functional enrichments demonstrate consistent patterns across xenarthran lineages. Next, we examined biological processes related to metabolism, revealing that retrocopies are strongly associated with key pathways essential for cellular function (Figure 4C). The most represented categories were carbohydrate metabolism, lipid metabolism, generation of precursor metabolites and energy, and amino acid metabolism, all of which were consistently observed across the five xenarthran species. Among the most prolific parental genes are ribosomal proteins (e.g., *RPS3A*, *RPL10*, *RPL4*, *RPL27A*), consistent with the biological processes to which these genes contribute (Figure 4D). A heatmap further shows distinct patterns of retrocopy accumulation across ribosomal protein families, with variation both among species and across gene families. Lineage-specific expansions are also present: ACP1 dominates in *C. didactylus* (>150 copies), GAPDH and ribosomal genes in *D. novemcinctus*, and PPIA (protein folding) in *T. tetradactyla*. In addition, *B. torquatus* and *C. hoffmanni* exhibit expansions of enzymes such as UBE2N, DHFR, and FDPS, which are involved in protein degradation, folate metabolism, and cholesterol/isoprenoid biosynthesis. Together, these patterns highlight a pronounced bias in retrotransposition toward parental genes with regulatory, enzymatic, and housekeeping functions.

### Retrocopy RNA-seq expression in *Choloepus didactylus* and *Dasypus novemcinctus*

Due to the high sequence similarity between retrocopies and their parental genes, especially in recently formed retrocopies, expression quantification is susceptible to read misassignment and false positives. To mitigate this, we implemented a simulation-based filtering strategy (see Methods) to exclude retrocopies that could not be reliably distinguished from parental transcripts. In addition, for highly similar, recently duplicated retrocopies, expression was aggregated across copies derived from the same parental gene. Using this approach, we quantified retrocopy expression in five tissues of *C. didactylus* (spleen, brain, liver, lung, and blood) and three tissues of *D. novemcinctus* (spleen, muscle, and liver) using RNA-seq data. A markedly higher proportion of retrocopies is expressed in *C. didactylus* than in *D. novemcinctus*: expression in at least one tissue was detected for 7,787 retrocopies (48% of retrocopies) in *C. didactylus*, compared to 3,615 retrocopies (27%) in *D. novemcinctus*. In both species, retrocopies consistently showed lower expression levels relative to their parental genes across all tissues (Supplementary Figure 4), consistent with retrocopy expression patterns described in other mammals^12,20^. Analysis of synonymous substitution rates (dS) from ORF–parental gene alignments indicated that expressed retrocopies span a broad range of insertion ages (Supplementary Figure 5). Notably, a significantly higher proportion of expressed retrocopies in *C. didactylus* had lower dS values relative to those in *D. novemcinctus* (Fisher’s exact test, dS < 0.2: p < 3.4 × 10⁻⁹, OR = 1.34, Supplementary Figure 5).

### In search of function: potentially domesticated retrocopies in *C. didactylus*

To investigate whether some retrocopies have been retained under functional constraint (i.e., domesticated), we focused on *C. didactylus*. We refer to retrocopy domestication as the evolutionary process by which initially non-functional retrocopy insertions acquire biological function and are subsequently maintained by purifying selection, as evidenced by reduced nonsynonymous substitution rates and sustained expression patterns. A common approach to infer functional constraint is to examine the ratio of nonsynonymous to synonymous substitutions (dN/dS) between retrocopies and their parental genes. However, for very recent insertions, high sequence identity can produce artificially low dN/dS values that primarily reflect recency rather than genuine purifying selection. The presence of retrocopies with orthologs only across sloths (i.e., originated somewhere in the branch leading to their last common ancestor ∼30 Mya) provides a valuable opportunity to overcome this bias and assess whether some retrocopies have been retained under constraint.

We defined candidate domesticated retrocopies as those that: (1) are shared only among sloths, (2) are expressed in at least one tissue, (3) encode an ORF at least 70% the length of the parental protein, and (4) have a dN/dS ratio below 0.5. Of 5123 sloth-orthologous retrocopies identified in *C. didactylus*, 2,347 show expression, 525 retain a long ORF, and 123 meet all four criteria. Manual inspection of alignments retained 52 high-confidence candidates. As a final quality control step, we further filtered the RNA-seq data to keep only retrocopies with at least one uniquely mapped read. Fourteen retrocopies were excluded because, although they showed signal in aggregated counts, they showed no uniquely mapped reads to the specific candidate loci (Supplementary Figure 6). This resulted in a final set of 38 retrocopies with strong evidence of potential domestication.

The 38 candidate retrocopies represent a core set of genes linked to metabolism, housekeeping, and stress response (Supplementary Table S1). Many derive from genes involved in mitochondrial, energy metabolism and lipid regulation (AUH, CHCHD4, CISD1, COX5B, PRPS2-like, MOSPD1, MRPS36, PLA2G12A), stress signaling and heat shock response (HSBP1, CARHSP1), ribosomal proteins (RPL, RPS), and RNA-binding and translation associated factors (SZRD1, TPRKB). Expression profiles (Figure 5A) revealed tissue-biased patterns, with many domesticated retrocopies related to mitochondrial and metabolic functions showing elevated expression in blood, brain, or liver. Stress-response retrocopies were preferentially expressed in blood, while ribosomal retrocopies tended to be either broadly expressed or enriched in specific tissues such as brain or lung. Functional annotation reinforced these categories, with GO terms highlighting roles in ribosome, organelle, catalytic activity, and stress response (Figure 5B). A protein–protein interaction (PPI) network (STRING, visualized in Cytoscape; Figure 5C) further underscored connections among mitochondrial, ribosomal, and RNA-binding parental genes.

**Figure 5.**
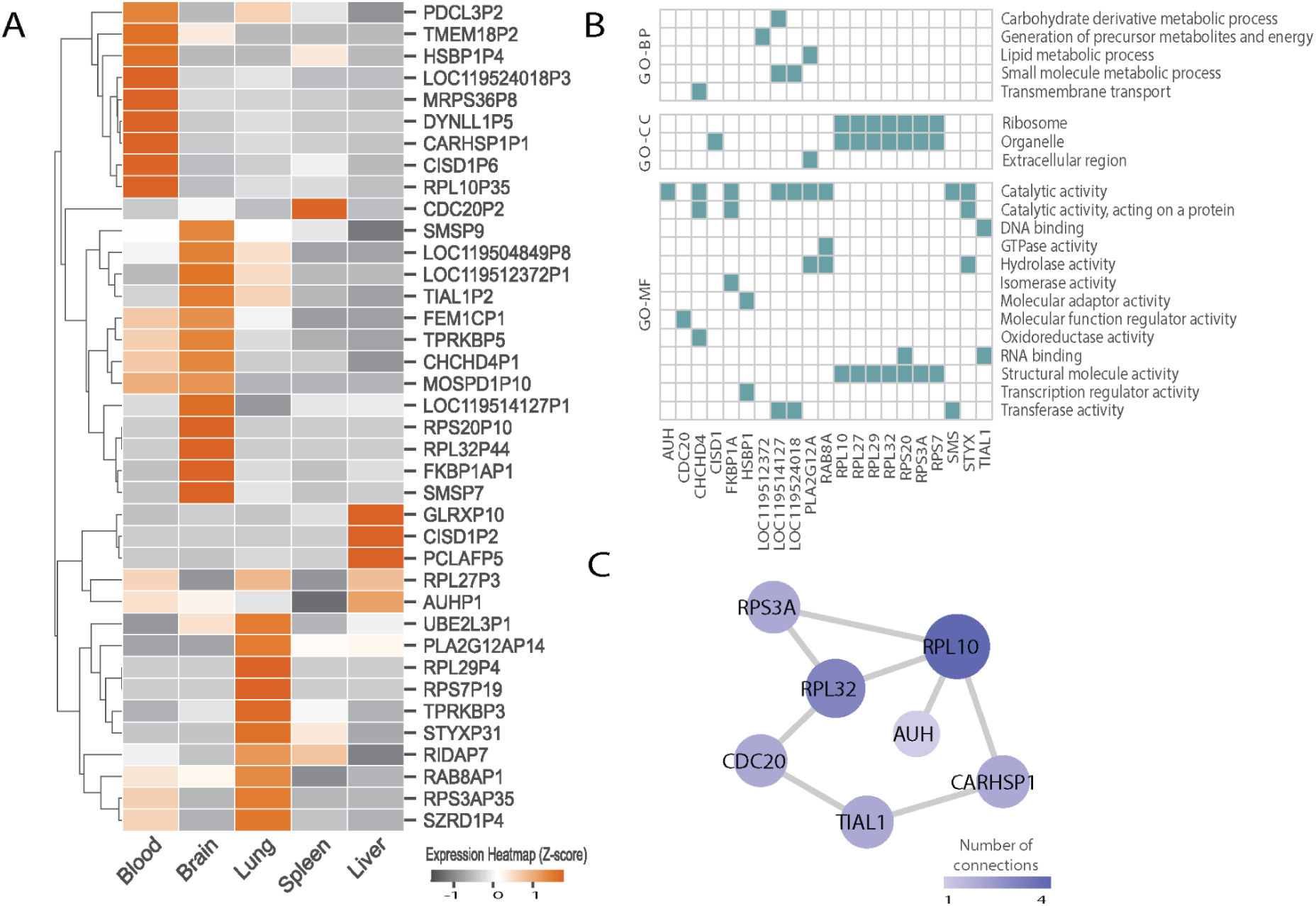
Expression profile and functional characterization of domesticated retrocopy candidates in *C. didactylus*. A) Heatmap of expression Z-scores normalized per retrocopy across five tissues. Rows represent the retrocopies’ names. Hierarchical clustering was performed using Euclidean distance and complete linkage to highlight expression patterns. B) Gene Ontology (GO) terms (BP: Biological Process, MF: Molecular Function, CC: Cellular Component) associated with the parental genes depicted in panel A. C) Protein-protein interaction (PPI) network illustrating interactions among the parental genes depicted in panel A, highlighting potential functional relationships.

## Discussion

### New high-quality genomes for Xenarthra

Here, we presented two new high-quality genome assemblies for Xenarthra, a placental mammalian clade long underrepresented in genomics. Previous available assemblies for Xenarthra, such as the Sanger-based genome of *Choloepus hoffmanni*, remain highly fragmented despite later Hi-C scaffolding (e.g., 117,974 gaps; contig N50 = 64 Kb). In contrast, our *C. didactylus* assembly reaches a contig N50 of 20 Mb, marking a substantial improvement for *Choloepus* species. Alongside improved references for *D. novemcinctus* and *B. torquatus*, the genomes of *C. didactylus* and *T. tetradactyla* presented here advance xenarthran genomics and provide a foundation for comparative studies, including the evolution and impact of retrocopies.

### A standardized annotation of retrocopy identification across mammalian genomes

Recently, some of us presented an updated version of RCPedia^19^, a comprehensive database cataloguing retrocopies across 44 mammalian genomes. RCPedia offers both an improved identification pipeline and a user-friendly interface for exploring retrocopy data. A significant challenge in retrocopy research is the variability in detection across studies, primarily due to methodological differences, including alignment criteria, filtering thresholds, and parental gene annotation strategies. Such inconsistencies often lead to large discrepancies in reported retrocopy numbers across species, underscoring the need for standardized pipelines in comparative analyses. Our identification pipeline focuses on biological hallmarks of retrotransposition, including tolerance for truncated insertions through a low size threshold, retention of 3′ mRNA sequences, and allowance for post-insertion TE activity within retrocopies. However, some limitations remain, such as the inability to detect mono-exonic parental genes, cases where the parental gene or reference transcript has been lost, or retrocopies extensively masked by subsequent TE insertions. To ensure robust cross-species comparisons, we applied our RCPedia pipeline to the newly assembled xenarthran genomes, revealing that Xenarthrans harbor the highest number of retrocopies reported for any animal clade to date.

### Repeats, retrocopy formation, and evolutionary dynamics

LINE1 activity has long been associated with retrocopy generation in humans and other mammals^11^, as the ORF2 protein preferentially binds poly-A tails, unlike other LINEs that recognize more complex motifs^21^. This association is also evident in Xenarthra, but the dynamics differ across lineages. Sloths, particularly *C. didactylus*, exhibit an enrichment of young, full-length, and expressed LINE1 families alongside a large number of recent retrocopies. In contrast, armadillo and anteater harbor older LINE1 repertoires, and their clade-specific retrocopies lack signatures of recent activity. Armadillos (Cingulata) are the most diverse xenarthran lineage (25 extant species), followed by sloths (Folivora, seven species) and anteaters (Vermilingua, eleven species)^3^. Our dataset includes both extant sloth genera (*Choloepus* and *Bradypus*), but only a single representative of armadillos and anteaters. Additional high-quality genomes for these lineages will therefore be critical to determine whether the observed retrocopy patterns reflect species-level processes or broader clade-level dynamics.

### Retrocopies as material for genomic innovation

The overall retrocopy landscape in Xenarhra mirrors patterns observed in other mammals^19^, with a strong enrichment for highly expressed and housekeeping genes. In *C. didactylus*, about half of retrocopies are transcriptionally active, compared with roughly one third in the armadillo. While a substantial portion of this expression may represent transcriptional noise, retrocopies have also been shown to modulate the expression of their parental genes, acting through mechanisms such as antisense interference, transcript competition, or regulatory disruption^22^. In addition, retrocopies can give rise to entirely novel genes through a process known as retrogene domestication, contributing new genetic material to host genomes^23^.

The presence of sloth-only orthologous retrocopies offered an opportunity to look for domesticated retrocopies in this lineage. Sloths are renowned for their exceptionally low metabolic rates—the lowest recorded among mammals—with basal metabolic rates reduced by 69–79% compared to expectations based on body mass^8^. Both *Bradypus* and *Choloepus* exhibit heterothermy, adjusting their internal body temperature in response to ambient temperature^9^, displaying a metabolic flexibility rarely seen in mammals. However, the link between this phenotype and the molecular biology of sloths is still not understood. Recent findings from some of us suggest relaxation in the mitochondrial genomes of both sloth genera (Johnson *et al.,* in preparation), potentially reflecting a greater tolerance for mutational accumulation due to their diminished energetic demand. In this study, we identified 38 retrocopies in *C. didactylus* with strong hallmarks of domestication. A substantial fraction of these retrocopies has parental genes functionally linked to mitochondrial processes and cellular metabolism. For example, AUH is a bifunctional mitochondrial protein involved in protein synthesis, RNA metabolism, biogenesis and overall mitochondrial function^24^; CHCHD4 is a central redox-sensitive import factor required for mitochondrial respiratory chain biogenesis^25^; CISD1 is a mitochondrial outer-membrane protein that maintains lipid homeostasis and regulates reactive oxygen species (ROS) production via Fe-S clusters redox reactions^26^; PLA2G12A participates in lipid metabolism^27^; GLRX is a crucial enzyme of the cell’s antioxidant defense system, and may buffer redox imbalance under conditions of compromised electron transport chain (ETC) efficiency; HSBP1 functions as a stress chaperone, potentially stabilizing protein folding under different metabolic profiles. Together, these patterns indicate strong involvement of mitochondrial and metabolic parental genes among domesticated retrocopies, pointing to an intriguing hypothesis: in the face of mitochondrial genome relaxation, domesticated retrocopies in sloths may act as compensatory buffers, sustaining essential bioenergetic and redox processes, or even providing adaptive routes for metabolic flexibility. Although this hypothesis remains to be experimentally validated—including whether molecular relaxation translates into impaired mitochondrial function in any cellular context—such compensatory roles could be particularly important for maintaining baseline metabolic function in an organism adapted to energy conservation. Further functional investigation of these candidates offers a promising direction for understanding genome evolution under conditions of low metabolic demand.

### Conclusions

Xenarthrans possess the highest number of gene duplications mediated by retrotransposition (retrocopies) reported for any mammalian lineage. Our results reveal lineage-specific differences in LINE1 activity and retrocopy formation: while activity appears to be older in armadillo and anteater, sloth genomes indicate ongoing LINE1 mobilization in cis (i.e., by retroduplication itself) and in trans (creating retrocopies), and therefore likely a continued impact on their genome architecture. Notably, we identify at least 38 sloth-specific retrocopies with strong signatures of domestication, several of which are functionally linked to metabolism. These may contribute to the distinctive phenotype observed in this highly specialized group of mammals.

## Methods

### Samples, sequencing and assembly

*Choloepus didactylus* (“Lama Su”) and *Tamandua tetradactyla* (“Anton”) were zoo specimens housed at Tierpark Berlin. Samples were collected post-euthanasia and immediately flash-frozen. For *C. didactylus*, spleen, blood, brain, lung, and liver were collected. High molecular weight DNA was extracted from the spleen for genome sequencing, and all tissues (including spleen) were used for RNA sequencing. For *T. tetradactyla*, spleen tissue was used for DNA sequencing only. All data is available on NCBI under Bioproject Accessions PRJNA678727 and PRJNA561940.

Genome assemblies for both species were generated using the Vertebrate Genomes Project (VGP) pipeline v1.6^28^. PacBio CLR reads were assembled with FALCON and FALCON-Unzip^29^, and haplotigs were removed using Purge-Dups^30^. The primary contigs were scaffolded using 10X Genomics linked reads, Bionano optical maps (Solve), and Hi-C data (Arima) with the SALSA2 pipeline^31^. Assembly polishing was performed in three rounds—first using Arrow with PacBio reads, followed by two rounds with FreeBayes and 10X data^32^. Final decontamination and manual curation were conducted following the methods described in Howe et al.^33^. Assemblies are available on NCBI under accessions GCF_015220235.1 and GCA_023851605.1.

### Genome annotations and other datasets

Short-read RNA sequencing of all *Choloepus didactylus* tissues was performed using a combination of ribosomal RNA depletion (blood) and poly(A) enrichment (spleen, brain, liver, lung). The resulting reads were submitted to NCBI (accession PRJNA516733) and used for genome annotation with the NCBI Eukaryotic Genome Annotation Pipeline. The annotated *C. didactylus* genome is publicly available on NCBI.

Three additional publicly available xenarthran genomes were included in our analyses: *Dasypus novemcinctus* (GCA_030445035.2), annotated via the NCBI pipeline; *Choloepus hoffmanni* (DNA Zoo^34,35^, accession ABVD00000000.2); and *Bradypus torquatus* (GCA_963992745.1). For *C. hoffmann*i and *B. torquatus*, we used TOGA annotations^36^ provided by the Hiller’s lab at the Senckenberg Institute and available at: https://genome.senckenberg.de/download/TOGA/human_hg38_reference/Xenarthra/.

### Repeat annotation, masking and identification of full-length LINE1 elements

Repeats in the genomes of *Choloepus didactylus, Choloepus hoffmanni, Bradypus torquatus, Tamandua tetradactyla,* and *Dasypus novemcinctus* were identified using EarlGrey^37^. The resulting species-specific repeat libraries were then combined into a single custom library. All five genomes were subsequently masked and annotated with RepeatMasker using this unified repeat library. Kimura two-parameter substitution levels between each repeat copy and its corresponding consensus sequence were estimated using the calcDivergenceFromAlign.pl script, from RepeatMasker v4.1.5. Repeat landscape plots were generated with custom python scripts.

To identify full-length LINE1 retrotransposons, we followed the approach described by Schartl et al. 2024^38^. LINE1 sequences were extracted based on their genomic coordinates, and open reading frames (ORFs) of at least 600 amino acids were identified using EMBOSS getorf. After further filtering, elements were classified as full-length if they contained two ORFs of ≥600 amino acids that spanned at least 90% of both the endonuclease and reverse transcriptase domains.

### Retrocopy identification Pipeline

Retrocopies were identified using our improved pipeline from RCPedia^19^. The approach is based on detecting sequence similarity between multi-exon protein-coding genes (parental genes) and intronless genomic alignments (retrocopies). Messenger RNA (mRNA) sequences were extracted from annotated transcript coordinates using the gffread algorithm^39^, and aligned to the genome with the LAST aligner (lastal -D1000)^40^ to identify candidate retrocopy insertions. Alignments were retained if they exceeded 120 base pairs in length. To prevent the inclusion of potential chromosomal duplications, candidates located more than 200,000 base pairs from their parental gene were removed. Retrocopies had to preserve at least one of the last three exon-exon boundaries of the parental gene, consistent with the expected reverse transcription mechanism from the poly-A tail. Additionally, alignments containing 40% or more repetitive elements, as identified by RepeatMasker, simpleRepeats, and windowMasker annotations, were excluded. A distance filter restricted retrocopy insertions from the same parental gene to at least 500,000 base pairs apart, reducing false positives without relying on predefined blacklists. Retrocopies overlapping three or more exons of annotated coding genes were excluded, and parental genes with more than five retrocopies overlapping genes from the same gene family were discarded. In cases where multiple alignments originated from the same parental gene, the best-scoring alignment was retained based on sequence identity and match percentage. When ties occurred, the alignment with the highest match/(match + mismatch) ratio was selected. Additionally, continuous alignments from the same mRNA transcript separated by up to 6,000 base pairs were merged, accounting for gaps introduced by repetitive elements.

### Retrocopy orthology inferences

To identify potential orthologous retrocopies, we extracted the genomic sequence of each retrocopy along with 3,000 base pairs of flanking sequence on both sides. Pairwise alignments were performed using LASTZ [Improved Pairwise Alignment of Genomic DNA, 2007] between each flanked retrocopy region and all corresponding retrocopy regions (including ±3,000 bp flanks) from other genomes, including Bonobo (*Pan paniscus*, GCF_013052645.1), Budgerigar (*Melopsittacus undulatus*, GCF_012275295.1), Cat (*Felis catus*, GCF_000181335.3), Chicken (*Gallus gallus*, GCF_000002315.6), Chimpanzee (*Pan troglodytes*, GCF_002880755.1), Chinese hamster (*Cricetulus griseus*, GCF_003668045.3), Cow (*Bos taurus*, GCF_002263795.1), Crab-eating macaque (*Macaca fascicularis*, GCF_000364345.1), Dog (*Canis lupus familiaris*, GCF_014441545.1), Dolphin (*Tursiops truncatus*, GCF_011762595.1), Drosophila (*Drosophila melanogaster*, GCF_000001215.4), Egyptian rousette (*Rousettus aegyptiacus*, GCF_014176215.1), Gibbon (*Nomascus leucogenys*, GCF_006542625.1), Golden snub-nosed monkey (*Rhinopithecus roxellana*, GCF_007565055.1), Gorilla (*Gorilla gorilla gorilla*, GCF_008122165.1), Greater horseshoe bat (*Rhinolophus ferrumequinum*, GCA_014108255.1), Greater mouse-eared bat (*Myotis myotis*, GCF_014108235.1), Green monkey (*Chlorocebus sabaeus*, GCF_000409795.2), Horse (*Equus caballus*, GCF_002863925.1), Human (*Homo sapiens*, GCF_000001405.39), Kuhl’s pipistrelle (*Pipistrellus kuhlii*, GCF_014108245.1), Lizard (*Anolis carolinensis*, GCF_000090745.1), Marmoset (*Callithrix jacchus*, GCF_009663435.1), Mouse (*Mus musculus*, GCF_000001635.27), Mouse lemur (*Microcebus murinus*, GCF_000165445.2), Opossum (*Monodelphis domestica*, GCF_000002295.2), Orangutan (*Pongo pygmaeus abelii*, GCF_002880775.1), Painted Turtle (*Chrysemys picta bellii*, GCF_000241765.4), Pale spear-nosed bat (*Phyllostomus discolor*, GCA_014049915.1), Panda (*Ailuropoda melanoleuca*, GCF_002007445.1), Pig (*Sus scrofa*, GCF_000003025.6), Platypus (*Ornithorhynchus anatinus*, GCF_004115215.1), Rabbit (*Oryctolagus cuniculus*, GCF_000003625.3), Rat (*Rattus norvegicus*, GCF_000001895.5), Rhesus (*Macaca mulatta*, GCF_003339765.1), Sheep (*Ovis aries*, GCF_002742125.1), Tasmanian Devil (*Sarcophilus harrisii*, GCF_902635505.1), Turkey (*Meleagris gallopavo*, GCF_000146605.3), Velvety free-tailed_bat (*Molossus molossus*, GCF_014108415.1), Zebra Finch (*Taeniopygia guttata*, GCF_008822105.2), Zebrafish (*Danio rerio*, GCF_000002035.6), as previously described^19^. Alignment coverage exceeding 50% and identity surpassing 60% were required. Furthermore, we ensured that at least 50% of the retrocopy aligned with the designated target region. When multiple possible orthologs were identified, the alignment with the highest coverage and identity was selected for each retrocopy. Retrocopy homology was classified into five broad categories: (1) species-specific, referring to retrocopies with no detectable homology in any other species analyzed; (2) sloth-shared, those homologous in two or all sloth species; (3) xenarthra-shared, those with homology in two or more xenarthran species, excluding sloth-shared retrocopies; (4) elephant-shared, referring to retrocopies homologous with at least one elephant species; and (5) other-species-shared, comprising retrocopies with homology in at least one non-xenarthran and non-elephant species.

### GO annotation

We used Interproscan to annotate GO terms for parental genes of retrocopies. Later, we matched those GOs with GOslim using goatools map_to_slim.py^41^. We used python scripts to parse and plot shared GO terms.

### Retrocopies and LINE1 Expression Quantification in *Choloepus didactylus* and *Dasypus novemcinctus*

To ensure accurate quantification of retrocopy expression despite their high sequence similarity to parental genes, we applied a stringent *in silico* filtering strategy. We simulated paired-end RNA-Seq data (60 million reads, HiSeq 150 profile) using SANDY^42^ v1.0 with protein-coding transcripts as input, explicitly excluding regions overlapping exonic retrocopies (SANDY parameters --sequencing-type paired-end --quality-profile hiseq_150 --number-of-reads 60000000 --jobs 20). Quantification was performed with kallisto v0.48.0 [https://pachterlab.github.io/kallisto/about] (-t 12 -b 100). Retrocopies with detectable expression in the simulations were deemed potentially confounding and excluded from downstream analysis.

For LINE1s, we simulated reads from consensus sequences in our repeat library. A kallisto index including both LINE1 consensus sequences and protein-coding transcripts was used for quantification. Only LINE1 subfamilies with a ratio of observed-to-expected simulated expression between 0.8 and 1.15 were retained.

Experimental RNA-Seq data were obtained from five tissues of *C. didactylus*—spleen, brain, liver, lung, and blood (SRR10066840–SRR10066845) and 3 tissues of *D. novemcinctus* (SRR32745552–SRR32745554)—and quantified using kallisto against a database index built from: (i) filtered retrocopies, (ii) simulation-vetted LINE1 consensus sequences, and (iii) RefSeq protein-coding transcripts (Annotation Release 100) excluding retrocopy overlaps.

To avoid ambiguous quantification of closely related retrocopies, particularly those derived from the same parental gene, Choloepus-specific retrocopies sharing a parental origin were grouped as single transcriptional units. Read counts for grouped retrocopies were aggregated using custom scripts. LINE-1 expression was measured at the subfamily level and normalized as transcripts per million (TPM).

To validate expression in candidates of domestication in *C. didactylus*, we complemented kallisto quantification with STAR^43^ v2.7.7a alignments (--outSAMtype BAM SortedByCoordinate --outSAMattributes NH HI AS nM MD XS), retaining only uniquely mapped reads. Reads aligning to parental genes or other recent duplicates were excluded. STAR-derived values were compared to kallisto estimates to assess consistency.

### dNdS estimation of retrocopies

To estimate nonsynonymous to synonymous substitution rate ratios (dN/dS), we first predicted open reading frames (ORFs) (minimal length=75aa) in retrocopies using TransDecoder. Predicted amino acid sequences for each retrocopy were aligned to its corresponding parental gene’s protein sequence using ClustalW. Codon-aware nucleotide alignments were then generated with PAL2NAL^44^, based on the protein alignments and underlying coding sequences. Synonymous (dS) and nonsynonymous (dN) substitution rates were calculated using codeml from the PAML package^45^, using the following parameters: CodonFreq = 2, model = 0, Nsites = 0, fix_omega = 0, and omega = 0.4.

### Domestication candidates selection

We parsed files from codon-aware alignments, codeML output and RNA-seq expression and defined candidate domesticated retrocopies as those that: (1) are shared among sloths, (2) are expressed in at least one tissue, (3) encode an ORF at least 70% the length of the parental protein, and (4) have a dN/dS ratio below 0.5.

### Protein-protein interaction analysis

We performed protein-protein interaction (PPI) analysis among parental genes of possibly domesticated retrocopies using Cytoscape (v. 3.10.3)^46^, based on interaction data from the STRING database (v. 11.5)^47^. Only interactions with a confidence score greater than 0.15 were retained.

## Data accessibility statement

All data for the new genomes is available on NCBI under Bioproject Accessions PRJNA678727 and PRJNA561940. Assemblies are available on NCBI under accessions GCF_015220235.1 and GCA_023851605.1. *C. didactylus* RNA-Seq reads can be found under the accession number PRJNA516733. Coordinates of retrocopy predictions can be found at RCPedia (https://www.rcpediadb.org/).

## Acknowledgements

We thank Jeffrey Padberg, Trygve Bakken, and Rebecca Hodge for allowing us to include the *Dasypus novemcinctus* genome in our analyses. We also thank Dr. Richard K. Wilson and The Genome Institute at Washington University School of Medicine for generating the *Choloepus hoffmanni* genome assembly used in this study. We are grateful to Gudrun Wibbelt for providing high-quality samples of the two-toed sloth “Lama Su” and the lesser anteater “Anton.” This work was supported by the European Union’s Horizon 2020 research and innovation programme under the Marie Skłodowska-Curie grant agreement No. 750747, and by the Wellcome Trust (Grant 220540/Z/20/A, Wellcome Sanger Institute Quinquennial Review 2021–2026). This work was also partially supported by the São Paulo Research Foundation (FAPESP) grant 2018/15579-8 awarded to PAFG, HBC, and RLVM.

## Authors contributions

M.U.S., C.J.M., H.B.C., and P.A.F.G. conceptualised the project. S.K. generated sequencing data. M.U.S. and S.M. assembled genomes. A.T. manually curated genomes. M.U.S., H.B.C., and R.L.V.M. analysed the data. G.D.A.G. performed GO enrichment, prepared figures; M.B., A.S., C.J.M., P.A.F.G. reviewed results. M.U.S., G.M., M.B., P.A.F.G., and C.J.M. acquired funding. M.B., P.A.F.G., and C.J.M. supervised the project. M.U.S. drafted the manuscript. All authors revised and approved the manuscript.

