## Supplementary Figures and Tables for "Retrocopy formation and domestication shape genome evolution in sloths and other xenarthrans"

### Supplementary Figure 1

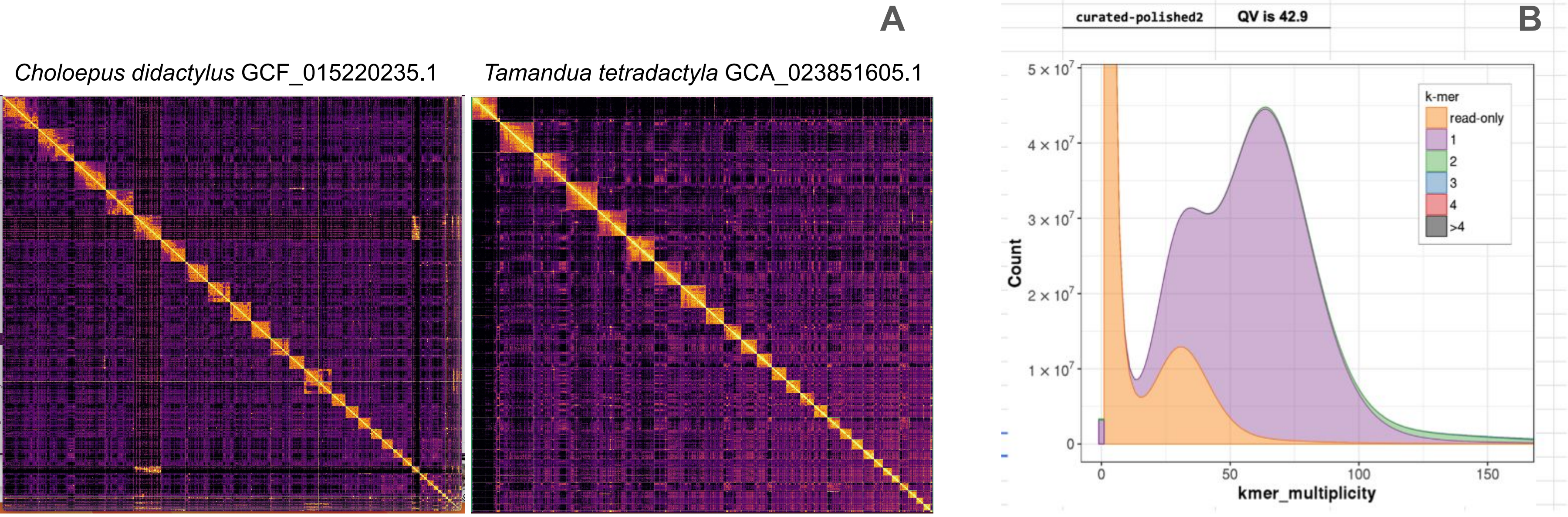


**Supplementary Figure 1|A:** Hi-C heatmaps for *Choloepus didactylus* and *Tamandua tetradactyla*. **B**: Merqury plot for *C. didactylus* final assembly: kmer counts show low levels of duplicated kmers in homozygous and heterozygous genome coverage peaks and distribution.

**Supplementary Table 1**: General metrics of publicly available Xenarthra genomes included in this study

| Species | *Choloepus hoffmanni* | *Bradypus torquatus* | *Dasypus novemcinctus* |
|---|---|---|---|
| Accession | DNAzoo C_hoffmanni-2.0.1_HiC.fasta.gz | GCA_963992745.1 | GCF_030445035.2 |
| Total length | 3,293,892,468 | 3,128,591,653 | 3,610,551,914 |
| Scaffold N50 | 140,950,122 | 156,600,181 | 127,081,865 |
| Contig N50 | 64,321 | 4,748,776 | 13,963,394 |
| Scaffold count* | 253,016 | 2,915 | 546 |
| BUSCO** mammalian_odb10 | C:94.2%[S:89.7%,D:4.5%],F:1.9%,M:3.9%,n:9226 | C:94.4%[S:90.1%,D:4.3%],F:1.1%,M:4.5%,n:9226 | C:93.8%[S:90.6%,D:3.2%],F:1.7%,M:4.5%,n:9226 |

*In cases where identified, Including the mitochondrial genome as a scaffold
**BUSCO: Benchmarking using single-copy orthologues

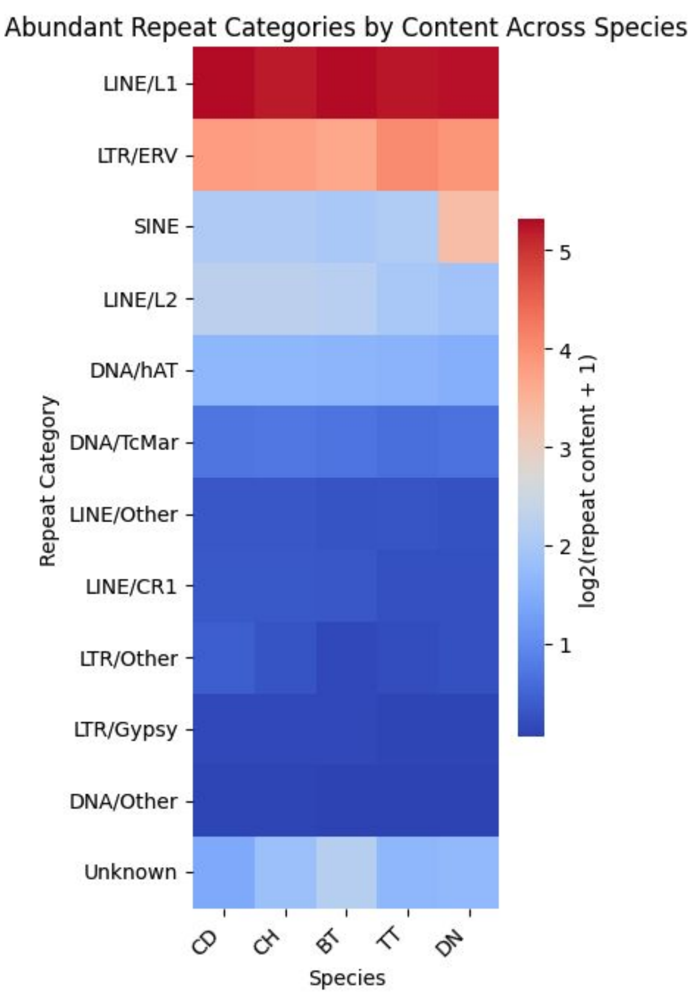


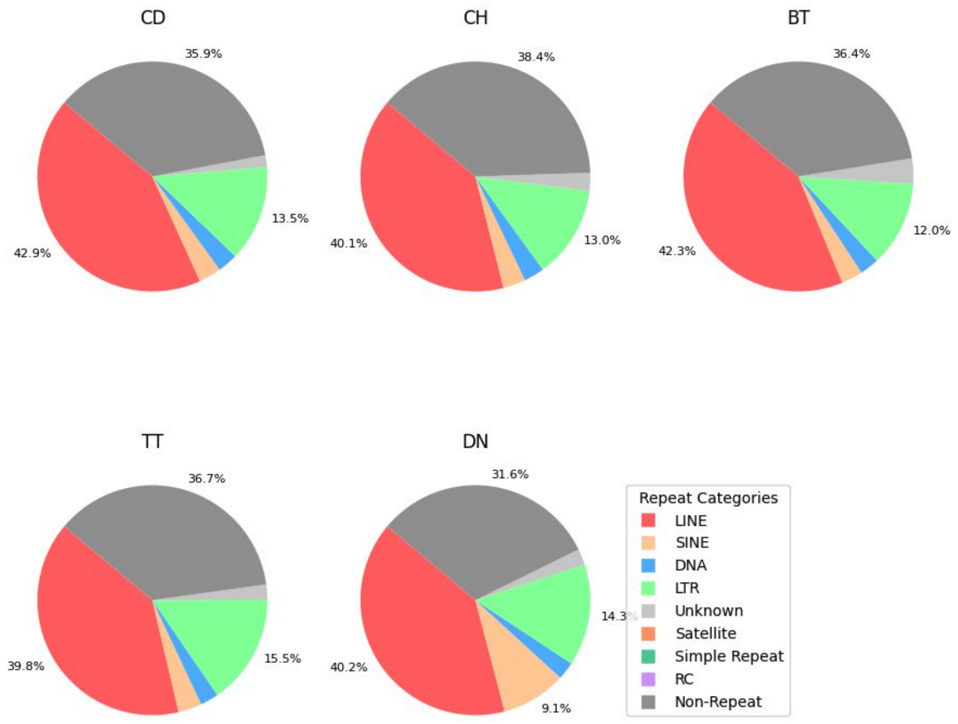


**Supplementary Figure 2**: General repeat content of Xenarthra genomes. Species abbreviations: CD, *C. didactylus*; CH, *C. hoffmanni*; BT, *B. torquatus*; TT, *T. tetradactyla*; DN, *D. novemcinctus*.

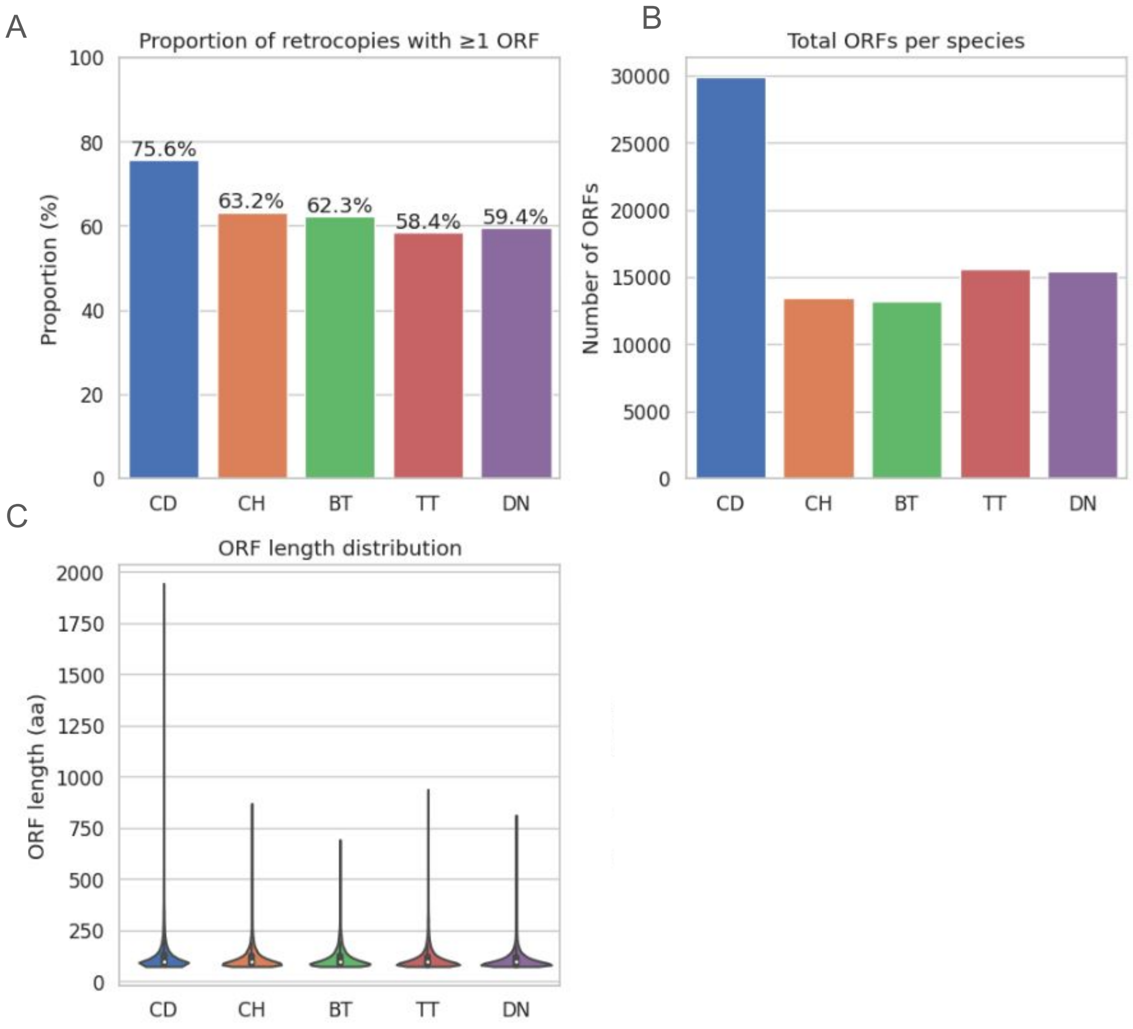


**Supplementary Figure 3**: Retrocopy-predicted ORFs. A) Proportion of retrocopies yielding ORFs, B) Total predicted ORFs per species, C) Length of retrocopy-predicted ORFs. Species abbreviations: CD, *C. didactylus*; CH, *C. hoffmanni*; BT, *B. torquatus*; TT, *T. tetradactyla*; DN, *D. novemcinctus*.

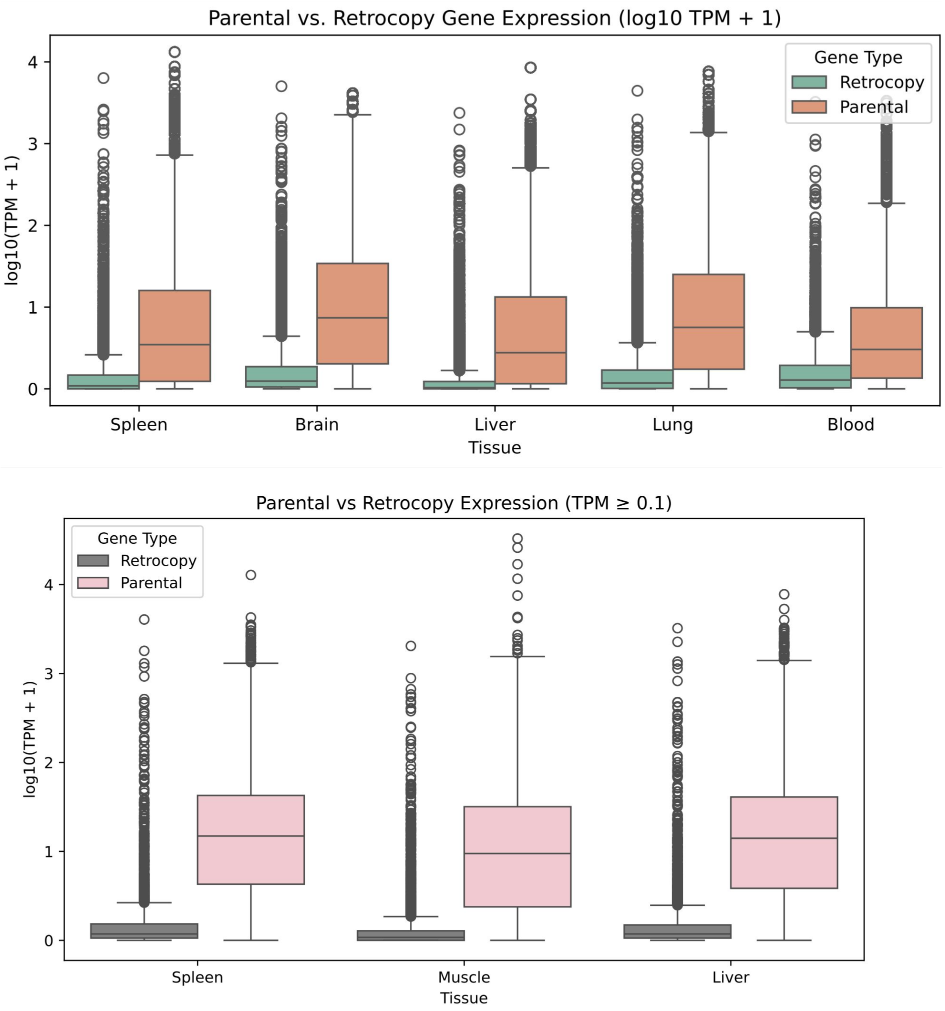


A

B

**Supplementary Figure 4**: RNA-seq parental genes and retrocopies expression in (A) *C. didactylus* and (B) *D. novemcinctus*

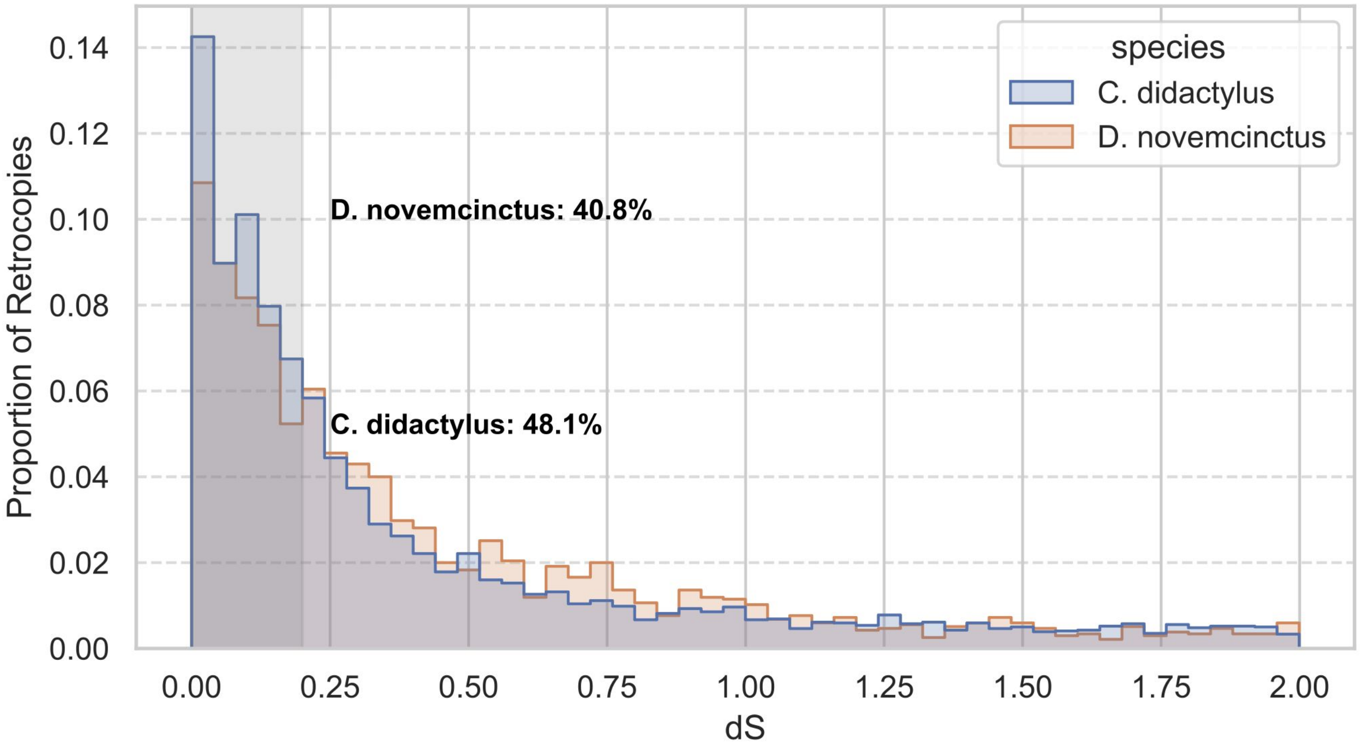


**Supplementary Figure 5** Distribution of synonymous substitution rates (dS) for expressed retrocopies in *C. didactylus* and *D. novemcinctus*. The overlay histogram shows the proportion of expressed retrocopies across dS bins, with the shaded region highlighting young retrocopies (dS < 0.2). A significantly higher proportion of expressed retrocopies in C. didactylus fall within this low-dS range compared to *D. novemcinctus* (Fisher's exact test, $p \approx 3.4 \times 10^{-9}$, odds ratio = 1.34).

**Supplementary Figure 6**: counts of uniquely mapped RNA-Seq reads in domesticated retrocopies of *C. didactylus*

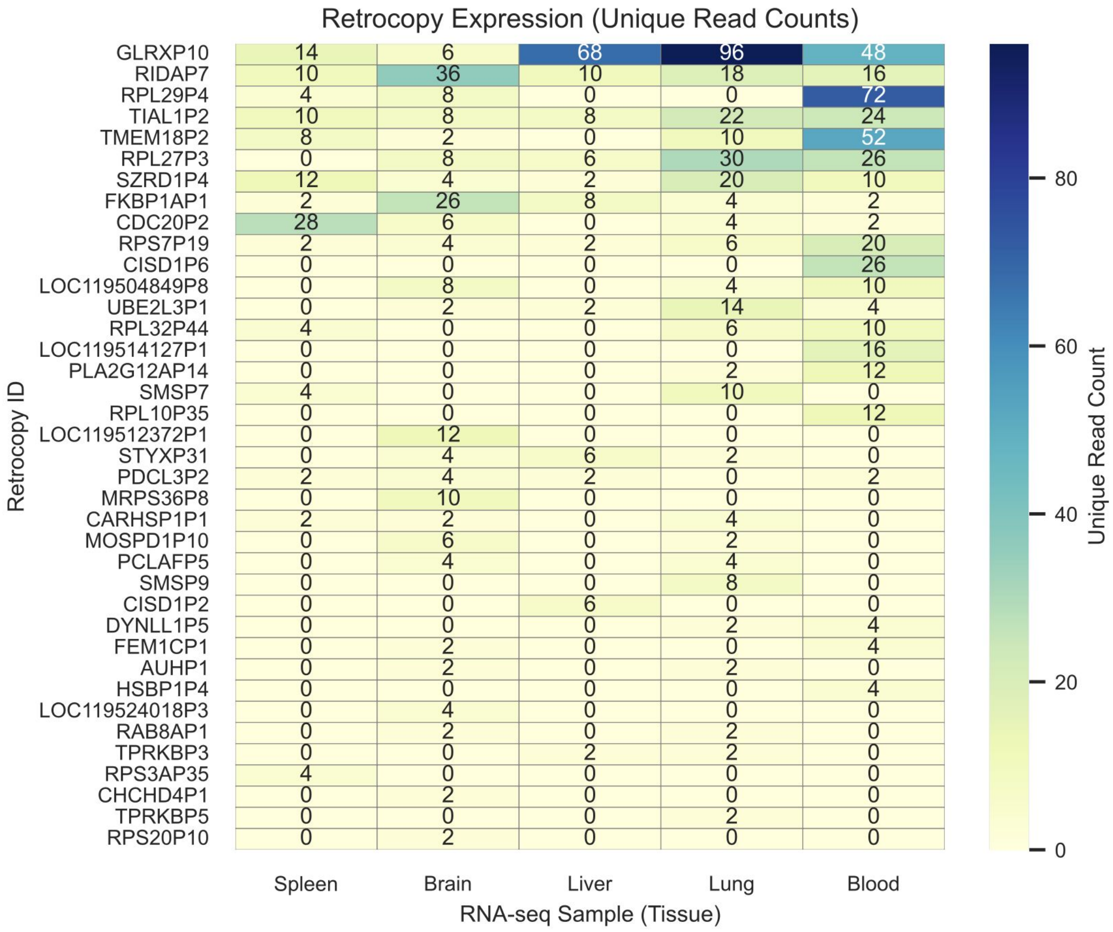

**Supplementary Table 2**: NCBI-assigned functions to parental genes given rise to domesticated retrocopies in *C. didactylus*

| symbol | name |
|---|---|
| AUH | AU RNA binding methylglutaconyl-CoA hydratase, transcript variant X2 |
| CARHSP1 | calcium regulated heat stable protein 1, transcript variant X3 |
| CDC20 | cell division cycle 20 |
| CHCHD4 | coiled-coil-helix-coiled-coil-helix domain containing 4 |
| CISD1 | CDGSH iron sulfur domain 1, transcript variant X2 |
| DYNLL1 | dynein light chain LC8-type 1, transcript variant X2 |
| FEM1C | fem-1 homolog C |
| FKBP1A | FKBP prolyl isomerase 1A |
| GLRX | glutaredoxin, transcript variant X3 |
| HSBP1 | heat shock factor binding protein 1 |
| LOC119504849 | guanine nucleotide-binding protein G(I)/G(S)/G(O) subunit gamma-10-like |
| LOC119512372 | cytochrome c oxidase subunit 5B, mitochondrial |
| LOC119514127 | nucleoside diphosphate kinase B, transcript variant X2 |
| LOC119524018 | ribose-phosphate pyrophosphokinase 2-like |
| MOSPD1 | motile sperm domain containing 1, transcript variant X2 |
| MRPS36 | mitochondrial ribosomal protein S36, transcript variant X2 |
| PCLAF | PCNA clamp associated factor, transcript variant X2 |
| PDCL3 | phosducin like 3, transcript variant X2 |
| PLA2G12A | phospholipase A2 group XIIA, transcript variant X1 |
| RAB8A | RAB8A, member RAS oncogene family |
| RIDA | reactive intermediate imine deaminase A homolog |
| RPL10 | ribosomal protein L10 |
| RPL27 | ribosomal protein L27 |
| RPL29 | ribosomal protein L29, transcript variant X1 |
| RPL32 | ribosomal protein L32 |
| RPS20 | ribosomal protein S20 |
| RPS3A | ribosomal protein S3A |
| RPS7 | ribosomal protein S7 |
| SMS | spermine synthase |
| STYX | serine/threonine/tyrosine interacting protein, transcript variant X1 |
| SZRD1 | SUZ RNA binding domain containing 1, transcript variant X4 |
| TIAL1 | TIA1 cytotoxic granule associated RNA binding protein like 1, transcript variant X7 |
| TMEM18 | transmembrane protein 18, transcript variant X1 |
| TPRKB | TP53RK binding protein, transcript variant X1 |
| UBE2L3 | ubiquitin conjugating enzyme E2 L3 |
